# Age-dependent lipid droplet-rich microglia worsen stroke outcome in old mice

**DOI:** 10.1101/2022.03.14.484305

**Authors:** Maria Arbaizar-Rovirosa, Mattia Gallizioli, Jordi Pedragosa, Juan J. Lozano, Carme Casal, Albert Pol, Anna M. Planas

## Abstract

Microglial cells of the aged brain manifest signs of dysfunction that could contribute to the worse neurological outcome of stroke in the elderly. Treatment with colony-stimulating factor-1 receptor antagonists enable transient microglia depletion that can be followed by microglia repopulation after interruption of the treatment, causing no known harm to mice. Using this strategy, we aimed to restore microglia function and ameliorate stroke outcome in aged mice. Cerebral ischemia/reperfusion induces strong innate immune responses in microglia highlighted by prominent type I interferon signaling, together with cellular metabolic perturbances and lipid droplet biogenesis in young mice. In aged mice, a subset of microglia accumulates lipid droplets under steady state and displays exacerbated innate immune responses after stroke. Microglia renewal in old mice reduces the lipid droplet content, prevents the ischemia-induced exaggerated type I interferon response, and improves the neurological outcome of stroke. This study shows that age-dependent lipid droplet-enriched microglia contribute to impair stroke outcome in old mice.

## Introduction

Age is the main non-modifiable stroke risk factor and stroke outcome is worse in the elderly. The higher vulnerability of the aged brain to ischemia has been attributed to differences in the mechanisms of ischemic brain injury between aged and young individuals^1^. Moreover, elderly patients with stroke show worse responses to reperfusion therapies^2^. Several lines of evidence support that exacerbated brain inflammation and altered immunological response underlie the worse neurological impairment in aged subjects after cerebral ischemia/reperfusion^3^. The brain of aged mice shows a differential transcriptional profile to that of the young^4^. The expression of genes encoding pro-inflammatory and innate immune response molecules is exaggerated in the elderly, whereas genes involved in axonal and synaptic structure and activity are downregulated^4–7^. Aging impairs microglia functions^8^ particularly affecting white-matter associated microglia^9,10^. Recent studies identified subsets of dysfunctional microglia in the aging brain characterized by increased oxidative stress, inflammatory profile, abnormal lipid accumulation^11^, and increased lysosomal storage^12^.

Microglial cells play critical roles in the central nervous system during brain development^13^ and in adulthood to maintain brain homeostasis via microglia-neuron interactions^14,15^. Microglia prevent excessive neuronal depolarization following excitotoxic insults^16^ and exert some protective effects in cerebral ischemia^17–19^. Microglia survival is dependent on colony-stimulating factor 1 receptor (CSF1R). In rodents, genetic strategies to prevent microglial *Csf1r* expression or treatment with pharmacological inhibitors of CSF1R causes microglia depletion^20^. Upon removal of the CSF1R inhibitor, microglia repopulate the brain and several lines of evidence support that repopulated microglia exert beneficial effects^21,22^. Proliferating and non-proliferating microglia with a distinct transcriptomic profile are detected in the adult mouse brain during repopulation following microglia depletion^23^. Notably, microglia repopulation in the aged brain reversed some age-induced changes in microglia gene expression, and reduced lysosome enlargement and lipofuscin accumulation^24^. In an experimental model of traumatic brain injury, repopulating microglia stimulated neurogenesis and promoted recovery mediated by IL-6 dependent neuroprotection^25^. We hypothesized that restoring the microglia phenotype in the elderly may improve the functional outcome of stroke. We investigated the microglia response to stroke in young (3-4-months) and aged (21-22-months) mice and studied whether microglia renewal improved stroke outcome in the elderly. Our work shows that drug-induced microglia repopulation in the elderly rejuvenates some phenotypic traits of old microglia and improves stroke outcome.

## RESULTS

### Cerebral ischemia/reperfusion changes the transcriptomic profile and function of microglia

Microglia undergo strong phenotypic alterations due to brain injury following stroke. In line with the need to remove dead cells and cell debris, microglia morphology became more reactive and displayed engulfing phagosomes (Fig. 1A). In the periphery of infarction, electron microscopy showed microglia surrounding swollen synaptic vesicles (Fig. 1B). We studied the transcriptomic profile of microglia obtained via fluorescence-activated cell sorting (FACS) from the brain of mice after an episode of transient ischemia induced by 45 min intraluminal middle cerebral artery occlusion (MCAo) followed by reperfusion (Fig. 1C), as we previously reported^26^. RNAseq analysis showed marked changes in the microglia transcriptomic profile four days post-ischemia versus controls (Fig. 1D). According to gene ontology (GO) pathways and gene set enrichment analysis (GSEA), ischemia upregulated innate immune pathways in microglia with prominent enrichment of genes regulating lipid storage and defense response to virus, particularly interferon (IFN)-α and IFN-β (Fig. 1E,F). Other biological processes highly enriched in ischemic microglia included terms related to phagocytosis, lysosomes, and cholesterol storage (Fig. 1G). Notably, some of the abovementioned ischemia-induced differentially expressed genes (DEGs) in microglia are typically reported in disease-associated microglia (DAM) under diverse neurodegenerative conditions^27–28^. Expression of the DAM genes was similarly upregulated or downregulated in ischemic microglia. Therefore, microglia acquire features of a DAM-like profile within hours/days of an acute ischemic insult (Fig. 1H).

**Fig. 1.**
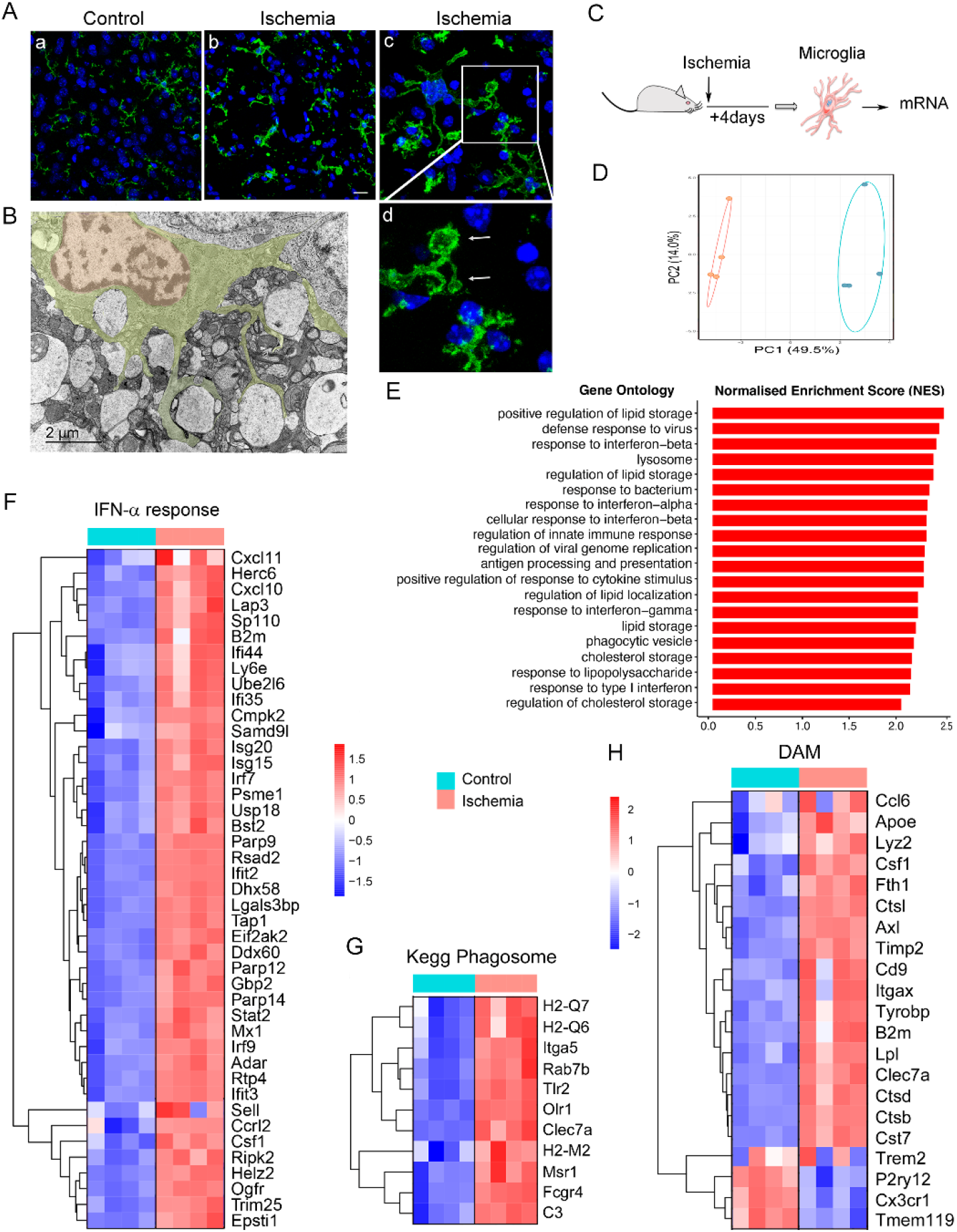
Transcriptomic and phenotypic changes in microglia after brain ischemia. A) Representative P2YR12 immunostaining (green) of microglia of wild type mice showing morphological differences between control microglia (a) and ischemic microglia (b-d). Ischemic microglial cells show typical phagocytic pouches (arrows). Nuclei are labeled with TO-PRO-3 (blue). Image (d) is a magnification of the area marked with a square in (c). Scale bar a, b: 20 μm; c: 10 μm; d: 4 μm. B) Transmission electron microscopy showing a microglial cell at the periphery of infarction one day post-ischemia surrounding remarkably swollen postsynaptic vesicles. Scale bar: 2 μm. C) Microglia were obtained using fluorescence-activated cell sorting (FACS) from the brain of control and ischemic young (3-4 months) male CX3CR1cre^ERT2^:Rosa26-tdT mice four days post-ischemia (n=4 mice per group). Microglia RNA was extracted for RNAseq analysis. D) Principal components analysis (PCA) shows sample distribution clearly separating microglia from control and ischemic mice. E) Gene Ontology analysis illustrating pathways enriched in microglia after brain ischemia. F) GSEA highlights the IFN-α response pathway as highly upregulated after ischemia. G) Genes of the Kegg pathway: *Phagosome* are upregulated in microglia after ischemia. Color scale as in (F). H) Most genes described as upregulated or downregulated in disease-associated microglia (DAM) are accordingly regulated in microglia four days post-ischemia.

### Ischemia induces the formation of lipid droplets in microglia

Conditions involving infection or inflammation in certain cell types cause accumulation of neutral lipids in the cytoplasm forming lipid droplets. Interestingly, lipid droplets are surrounded by a monolayer of phospholipids decorated with innate immune molecules, particularly proteins regulated by type I IFN that play a critical role in immune defence^29,30^. Given the conspicuous type I IFN response induced by cerebral ischemia in microglial cells, as well as the accompanying inflammatory response and enrichment in genes regulating lipid storage, we hypothesized that ischemia could induce lipid droplet biogenesis in microglia. We checked for ischemia-induced DEGs encoding typical lipid droplet membrane protein components, such as Plin1-5 family, and the reported IFN-induced lipid droplet-associated proteins.^29^ Brain ischemia in young mice increased at day four the microglial mRNA expression of Adipophilin (*Plin2*); Perilipin (*Plin3*); Hypoxia Inducible Lipid Droplet Associated (*Hilpda*); granulin precursor (*Grn*); Abhydrolase Domain Containing 5, Lysophosphatidic Acid Acyltransferase (*Abhd5*); Synaptosome Associated Protein 23 (*Snap23*); and lipoprotein lipase (*Lpl*) (Fig. 2A). Moreover, we detected ischemia-induced microglia mRNA upregulation of IFN-induced lipid droplet-associated molecules such as ISG15 Ubiquitin Like Modifier (*Isg15*); Ubiquitin Conjugating Enzyme E2 L6 (*Ube2L6*); Ubiquitin Specific Peptidase 18 (*Usp18*); Viperin (*Rsad2*); Ring Finger Protein 213 (*Rnf213*); Immunity Related GTPase M (*Irgm1*); and Interferon-Inducible GTPase 1 (*Iigp1*) (Fig. 2A). We also validated the ischemia-induced increased brain expression of Plin2 at the protein level by Western blotting (Fig. 2B). We identified the presence of lipid droplets in microglia by transmission electron microscopy of brain tissue 24h after ischemia (Fig. 2C). Furthermore, we observed lipid droplets in contact with lysosomes, supporting the occurrence of macrolipophagy as a mechanism that could supply energy to the cells. The presence of lipid droplets in ischemic microglia was quantified by flow cytometry after staining the cells with fluorescent Bodipy. While the percentage (mean±SD) of Bodipy^+^ CD45^low^CD11b^+^ microglia was 4.4±2.1% in the control brain, four days post-ischemia Bodipy^+^ microglia increased to 23.5±8.1% of total microglial cells in the ischemic brain hemisphere (Fig. 2D, E).

**Fig. 2.**
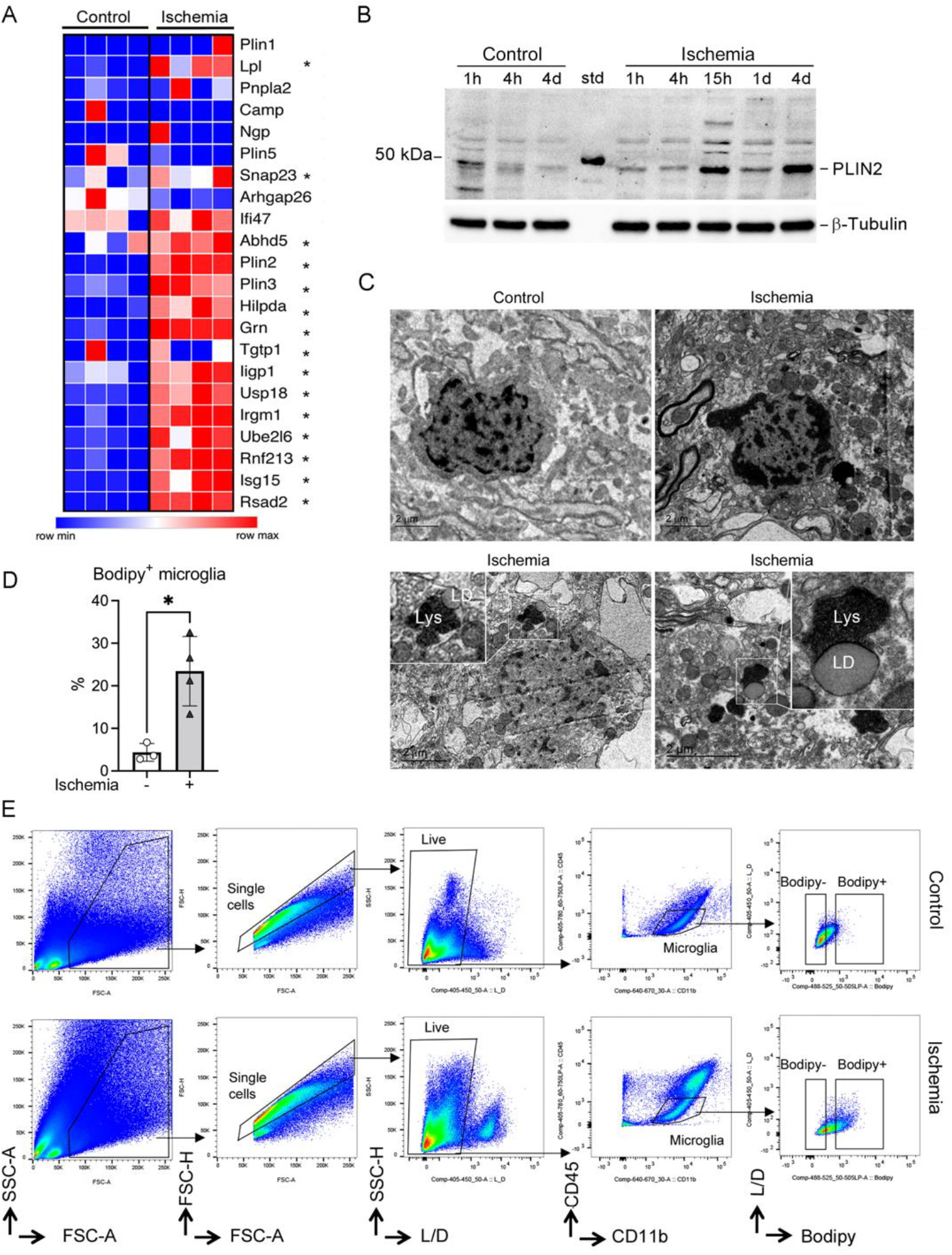
Microglia of young mice accumulate lipid droplets after brain ischemia. A) Expression of a selection of genes encoding lipid droplet-associated proteins in microglia sorted from control (n=4) and ischemic male mice (n=4) at day four post-ischemia (obtained from RNAseq data shown in Fig. 1). Heatmap illustrates ischemia-induced upregulation of genes marked with * indicating adjusted p value <0.001. B) Plin2 protein expression, as assessed by Western blotting in brain tissue obtained from the control (contralateral) and ischemic (ipsilateral) brain hemispheres at different time points post-ischemia, shows marked upregulation of Plin2 expression from 15h to four days post-ischemia. β-tubulin is the protein loading control. The ‘Std’ lane indicates the molecular weight standard. C) Transmission electron microscopy showing microglial cells in control or ischemic tissue one-day post-ischemia. Ischemia induces the formation of lipid droplets (LD) in microglia. LD are often seen near lysosomes (Lys). Insets in the lower panels are magnifications of the regions marked with a square. Scale bar 2 μm. D-E) Quantification of the percentage of CD11b^+^CD45^low^ microglia containing lipid droplets as Bodipy^+^ microglia using flow cytometry. While the presence of Bodipy^+^ microglia is negligible in control brain tissue (n=3), about 25% of the microglia becomes Bodipy^+^ four days post-ischemia (n=4) (* p=0.0117, unpaired *t*-test). Values correspond to individual mice and the mean±SD. (D) The flow cytometry gates to identify Bodipy^+^ microglia are shown in (E) for control and ischemic brain tissue. Gates were set based on fluorescence minus one (FMO) intensity values.

### Old mice show worse neurological outcome than young mice and microglia of old mice show exacerbated innate immune responses after stroke

Stroke caused more severe neurological deficits in old mice (21-22 month) than in young mice (3-4 months) and the effect was not attributable to larger lesion volume in the former (Fig. 3A). Comparison of the transcriptomic profile of microglia FACS-sorted from the brain four days post-ischemia in young and old mice showed 1,980 DEGs. Of these, 1,153 genes (58%) were upregulated in old ischemic mice, and so were most GSEA pathways indicating gain of function associated with age (Suppl. Table S1). Compared to microglia of young ischemic mice, microglia of old ischemic mice showed enrichment of GO pathways related to the innate immune system and antigen-presentation, amongst others (Fig. 3B). Enrichment of innate immune system and inflammatory pathways was highlighted by the GO terms *‘Response to interferon-beta’* (Fig. 3C)*, ‘Response to interferon-gamma’, ‘Response to bacterium’* and *‘Response to fungus’* (Suppl. Fig. S1). Many of these responses were already induced by ischemia in microglia of young mice (Fig. 1E), but they were upregulated further in microglia of old mice. Inflammatory responses were accompanied by upregulation of genes involved in the *‘Regulation of necroptotic cell death’* (Supp. Fig. S2A), and microglia of ischemic old mice also showed a prominent enrichment of antigen-presenting machinery (Suppl. Fig. S2B).

**Fig. 3.**
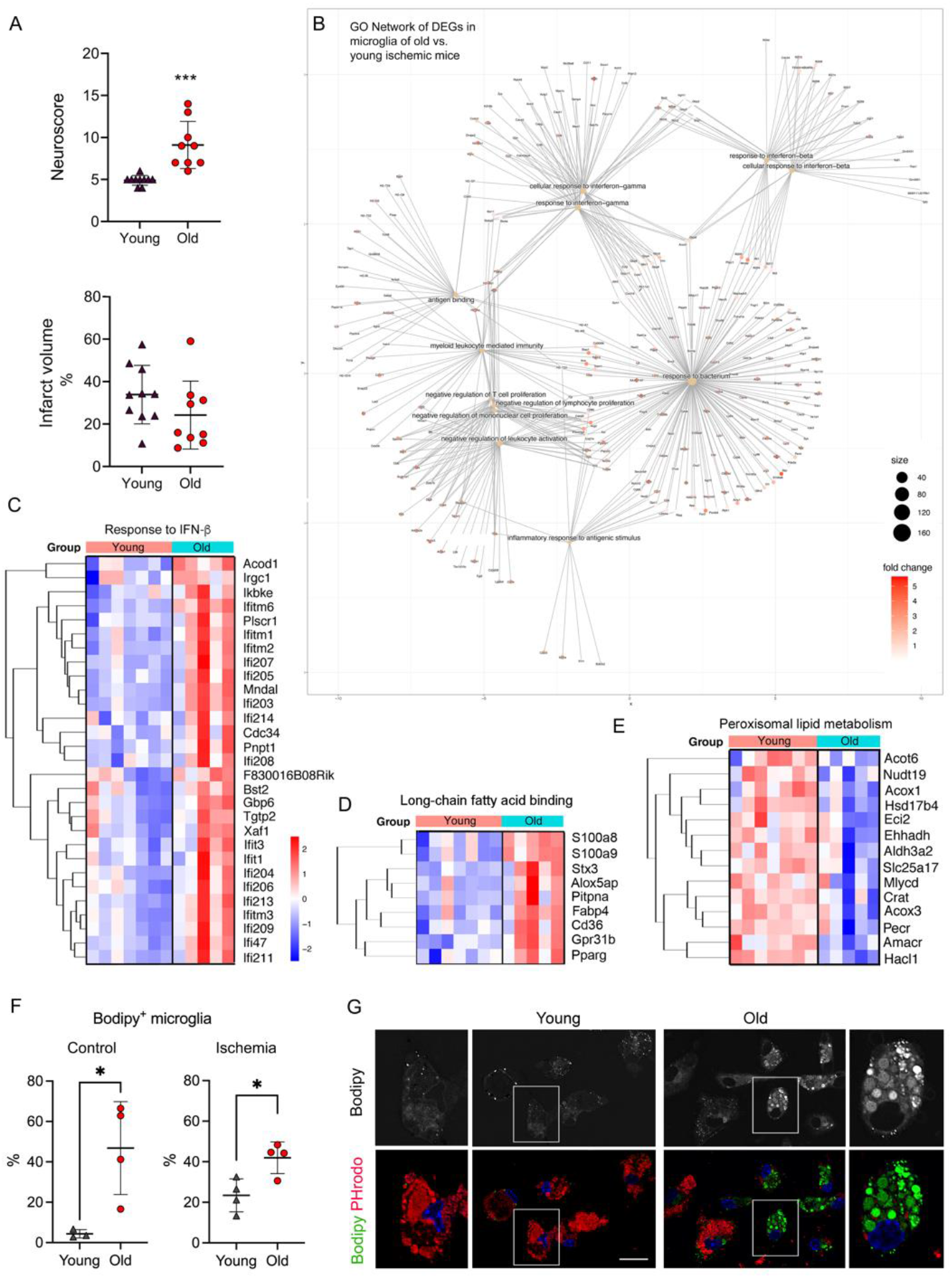
Worse stroke outcome in old mice and corresponding transcriptional profile of microglia. A) Neurological score and infarct volume four days post-ischemia in young (3-4 month) (n=10) and old (21-22 month) (n=9) female C57BL/6J mice. The neurological score was higher (worse) in old mice (Mann-Whitney test, p<0.0001), whereas there were no differences in infarct volume. B) Microglia mRNA was obtained from young (3-4 month; n=7) and old (21-22 month; n=5) female mice 4 days post-ischemia and studied by RNAseq. Cnetplot illustrates the network of genes enriched after ischemia in old vs. young mice linked to GO terms mainly related to innate immunity, inflammatory responses, and antigen presentation. C) Heatmaps of genes of representative innate immunity pathways enriched in microglia of old vs. young ischemic mice. D, E) Upregulation of genes related to long-chain fatty acid binding and downregulation of genes related to peroxisomal long-chain fatty acid metabolism in microglia of old mice. F) Microglia obtained from young and old male mice (n=3-4) 4 days post-ischemia (ipsilateral hemisphere) and control (contralateral) was studied by flow cytometry. Data from young and old mice were obtained in parallel, but for presentation purposes data of young mice are also shown in Fig. 2D. Old mice showed higher proportion of microglia containing lipid droplets in control (p=0.026) and ischemia (p=0.017) (*t*-test). Values show the mean±SD. G) Microglia from adult young and old mice using CD11b^+^ beads were kept in culture for 7 days and then exposed to red fluorescent pHrodo *E.coli* bioparticles and stained with Bodipy. Bodipy^+^ lipid droplets (white in the upper panels -raw intensity- and green in the corresponding lower panels) are clearly seen in microglia from old mice, but not young mice. The images in the bottom also show phagocytosed red bioparticles, which are hardly seen in lipid droplets containing cells. The square in each image in the center is magnified in the images on the left and right sides for young and old microglia, respectively. Scale bar= 20 μm.

In addition, microglia of old ischemic mice was enriched in pathways affecting *‘Phagocytosis’* (Supp. Fig. S2C), and processes involving lipids, e.g. *‘Long-chain fatty acid binding’* (Fig. 3D)*, ‘Positive regulation of fatty acid metabolic process’, ‘Cellular response to lipid’, ‘Lipid transport’*. In contrast, *‘Peroxisomal transport’, ‘Peroxisomal lipid metabolism’* and *‘Beta-oxidation of very long chain fatty acids’* pathways were downregulated in microglia of old ischemic mice (Fig. 3E; Supp. Fig. S2D). Deficiency in peroxisomal β-oxidation causes very long chain fatty acid accumulation as reported not only in the liver of aged subjects^32^ but also in the brain of Alzheimer’s disease patients^33^. Therefore, fatty acid disposal appears to be impaired in microglia of aged mice versus microglia of young mice after stroke.

These findings agree with the reported accumulation of lipid droplets in microglia of old mice under steady state, and the associated impairment of microglia phagocytic function.^11^ By flow cytometry we found that about 40% of microglial cells of old mice contained lipid droplets (Bodipy^+^) whereas there were negligible amounts in young mice (Fig. 3F). Again, after ischemia, old mice had more microglia cells containing lipid droplets than young mice (Fig. 3F). Interestingly, in old mice, ischemia did not increase the % of microglia with lipid droplets, in contrast to microglia of the young mice (Fig. 3F). Therefore, Bodipy^neg^ microglial cells of old mice do not generate lipid droplets in response to brain ischemia, suggesting that the old mice have a stable population of Bodipy^+^ microglia. We then obtained microglia from the brain of young and old mice by immunomagnetic isolation and maintained the cells in culture for 7 days. We analyzed the cells by confocal microscopy after exposure to red pHrodo *E. coli* bioparticles followed by Bodipy staining (Fig. 3G). A prominent subset of Bodipy^+^ microglia was only detected in the cultures obtained from old mice. Therefore, the culturing conditions did not remove the lipid droplets accumulated in microglia of old mice. Moreover, the microglia cells most enriched in lipid droplets did not contain or contained few phagocytosed bioparticles (Fig. 3G), indicating that they display impaired phagocytosis.

### Microglia repopulation restores certain transcriptional features of old microglia after ischemia

We then hypothesized that microglia renewal by depletion/repopulation could rejuvenate some features of the microglial phenotype in old mice, thus improving stroke outcome. Microglia can be depleted via several strategies^20,33^. We used treatment with the CSF1R antagonist PLX5622 provided in the diet as it strongly reduces brain microglia content after 2-3 weeks of treatment, as previously reported^18,21^. Microglia repopulate the brain after interruption of the PLX5622 diet and switch to control chow^21,34^. New repopulated microglia originates from brain-resident cells rather than peripheral hematopoietic cells, as demonstrated in chimeric mice generated by transplanting bone marrow from DsRed mice to recipient wild type mice that were later subjected to microglia depletion with PLX5622 and repopulation for seven days (Supp. Fig. S3A), in agreement with previous findings^34^. We treated young (3-4 months) and old (21-22 months) female mice with PLX5622 in the diet for three weeks and allowed microglia repopulation for different time periods: 3, 7, or 21 days. We then induced brain ischemia and studied the mice four days later (Supp. Fig. S3B). As expected, the PLX5622 diet reduced microglia cell number and the return to control diet increased microglia cell number, as seen in ischemic mice (Supp. Fig. S3C). For the subsequent experiments, we chose to induce ischemia after seven days of repopulation since, at this time point, microglia cell numbers had achieved values comparable to those of control mice and we could recover a substantial number of microglial cells for further study (Supp. Fig. S3C, D). We also checked whether leukocyte infiltration four days post-ischemia differed between old mice with or without microglia repopulation; we did not detect differences between these groups (Supp. Fig. S3E).

The comparison of the transcriptomic profile of renewed microglia versus original microglia of old mice after ischemia (Fig. 4A) detected 1,069 DEGs. Of these, 424 genes were upregulated and 645 genes were downregulated in renewed microglia. The downregulated GO terms (Suppl. Table S2) included innate immune responses like *‘Response to virus’* (Fig. 4B, C) and *‘Response to type I interferon’* (Supp. Fig. S4A). The GO terms *‘Pyroptosis’* (Suppl. Fig. S4B) and *‘Inflammasome complex’* were downregulated in repopulated microglia. Overall, the results showed that microglia repopulation in old mice reduced the innate immune and inflammatory responses displayed by these cells after ischemia. The antigen presentation capacity was also downregulated after microglia renewal, as illustrated by underrepresentation of the GO terms *‘Antigen binding’, ‘Antigen processing and presentation of endogenous antigen’*, and *‘Antigen processing and presentation of endogenous peptide antigen via MHC class I’* (e.g., Suppl. Fig. S4C). In contrast, renewed microglia of old ischemic mice displayed upregulation of pathways related to protein synthesis and metabolic activation (Supp. Table S2). We illustrate this finding by showing the upregulated genes of the GO term *‘Mitochondrial respirasome’* (Supp. Fig. S4D). Microglia renewal also seemed to affect lipid metabolism (Suppl. Fig. S4E), as shown by upregulation of the GSEA pathways *‘Fatty acid metabolism’* and *‘Metabolism of lipids’* highlighted by increased expression of genes involved in metabolizing very long chain fatty acids and peroxisome β-oxidation, cholesterol intracellular transport, and regulation of cholesterol biosynthetic route, amongst others. Overall, the transcriptomic profile of renewed microglia of old ischemic mice shows reduction of innate immune responses together with improvement of mitochondrial function and protein and lipid metabolism when compared with the original resident microglia of old ischemic mice.

**Fig. 4.**
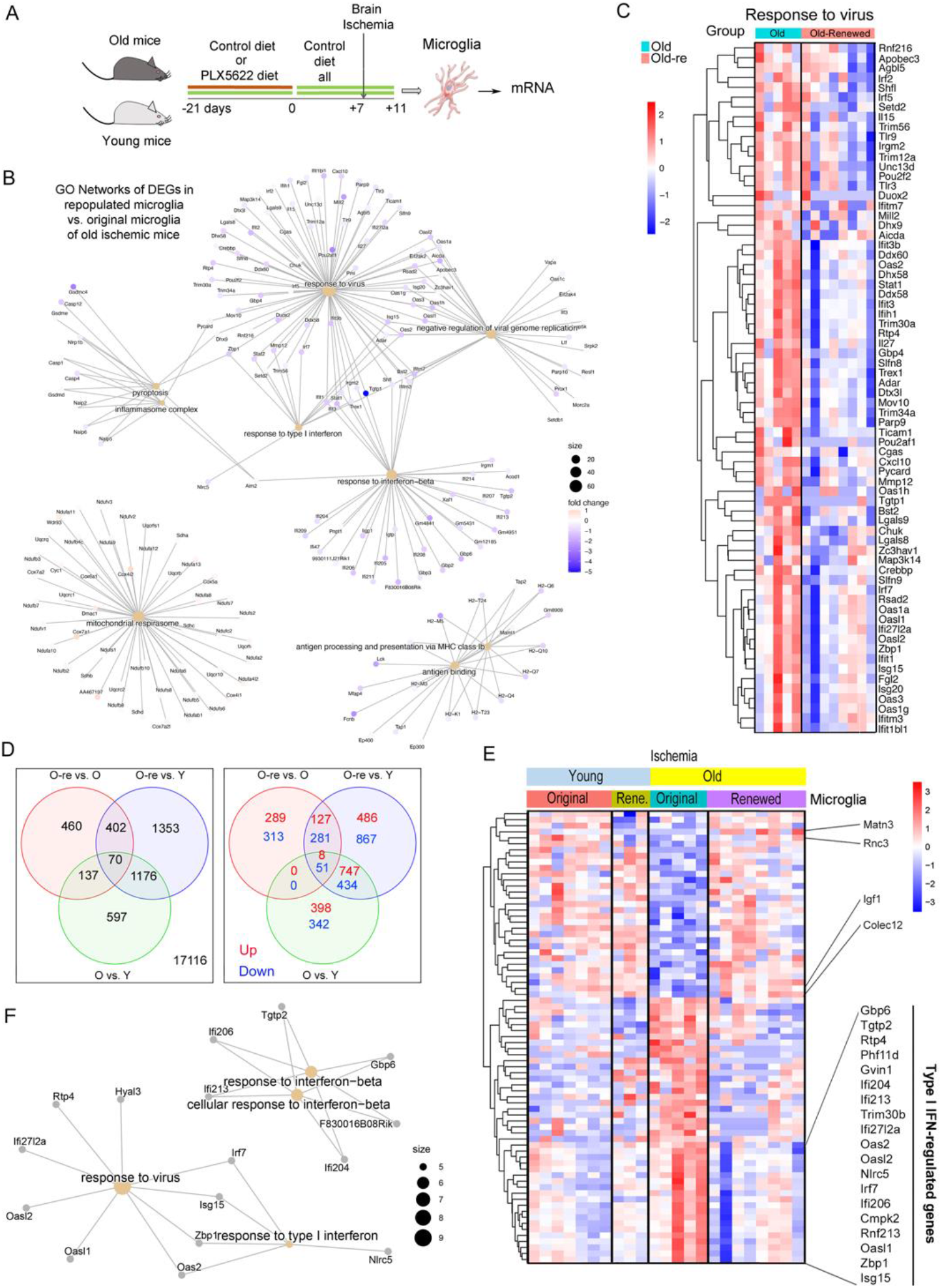
Microglia repopulation restores certain transcriptional features of old microglia after ischemia. A) Young (3-4 month) and old (21-22 month) female mice were fed control diet (n=12; n=5 old mice and n=7 young mice) or were treated with diet containing PLX5622 for 21 days to deplete microglia followed by restoration of control diet for seven days to induce microglia repopulation (n=11; n=8 old mice and n=3 young mice). Microglia was FACS sorted and RNA extracted for comparison of the transcriptomic profile of microglia obtained from old female ischemic mice with repopulated microglia. B) Cnetplot illustrating GO networks of DEGs in renewed microglia vs. original microglia of old mice after brain ischemia. C) Heatmap showing reduced expression of genes related to the GO term ‘Response to virus’ in renewed microglia. D) Venn diagram showing DEGs common and unique to the comparisons of the microglia obtained under the various experimental conditions after ischemia in female mice. There are 137 DEGs common for ‘microglia of old vs. young ischemic mice’ and ‘repopulated (re) vs. original microglia of old ischemic mice’ (upper diagram). Notably, none of these 137 genes is commonly upregulated or downregulated, signifying that they go in opposite directions in each comparison. E) Heatmap illustrating expression of 75 annotated genes of this group. Microglia renewal prevents the effect of old age by upregulating the expression of genes previously downregulated in old microglia and *vice versa*. F) Cnetplot illustrating the results of pathway analysis of the aforementioned 75 genes shows that the most enriched pathways correspond to IFN signaling.

For comparative purposes we also evaluated whether microglia renewal affected the transcriptomic profile in young ischemic mice (Supp. Table S3). In contrast to renewed microglia of old mice, renewed microglia of young mice did not downregulate the expression of innate immune response pathways, nor did it upregulate genes involved in mitochondrial respiration and lipid metabolism, whereas upregulation of genes involved in protein synthesis seemed to be common to renewed microglia regardless of the mouse age. Downregulated pathways in renewed microglia were mainly related to cell cycle, suggesting a tight cell cycle control at this stage of microglia repopulation.

### Microglia renewal in old mice prevents the exaggerated type I interferon innate immune response of old microglia to brain ischemia

We then examined the DEGs found when comparing the different groups of microglia of ischemic mice (age effect and microglia renewal effect in old mice) (Fig. 4D). We focused on the 137 DEGs common to ‘old vs. young’ (age effect) and ‘old renewed vs. old’ (microglia renewal effect in old mice), as illustrated in the Venn diagram (Fig. 4D, left; Supp. Table S4). Of note, none of these genes were commonly upregulated or downregulated in both comparisons (Fig. 4D, right). Therefore, these genes had to be regulated in opposite directions. Indeed, this was the case, as illustrated by the 75 annotated genes in the heatmap (Fig. 4E) showing either upregulation or downregulation of genes in microglia of old mice vs. microglia of young mice, and the restoration of gene expression by microglia renewal in old mice to the level of microglia of young mice. Among the genes downregulated in old microglia and upregulated after microglia renewal in old mice we found insulin growth factor 1 (*Igf1*), encoding a protein secreted by microglia with important neurogenic functions^35^. Within the genes upregulated in old microglia and restored by microglia renewal, the genes regulated by type I IFN stood out, e.g., *Isg15, Zbp1, Rnf213, Ifi206, and Irf7*, to mention but a few genes in this pathway. Other innate immune genes also downregulated by microglia renewal in old mice include the pattern recognition receptors *Cd209b* and *Fcnb*. We also identified restoration of expression levels of genes involved in lipid homeostasis and endoplasmic reticulum stress. Pathway analysis of these DEGs mainly detected type I IFN antiviral innate immune responses (Fig. 4F, Suppl. Table S5). Overall, these analyses reinforced the finding that repopulated microglia show less prominent innate immune responses compared with original microglia of ischemic old mice and suggest improvement of the cellular metabolic profile, particularly affecting lipid metabolism.

### Microglia renewal strongly reduces the age-dependent subset of microglia containing lipid droplets

Microglia of old ischemic mice upregulated the expression of some of the genes encoding lipid droplet-associated molecules even more than microglia of young ischemic mice (Fig. 5A). Importantly, microglia renewal in old mice significantly reduced the expression of type I IFN genes encoding for molecules associated with lipid droplets, e.g., *Rsad2, Isg15, Rnf213*, and *Irgm1* (Fig. 5B). Importantly, compared with untreated old mice, old mice with renewed microglia showed a reduced % of Bodipy^+^ microglia both under control and ischemic conditions, approaching levels like those observed in microglia of young ischemic mice, as assessed by flow cytometry (Fig. 5C, D). We then investigated whether microglia isolated from the brain of old mice and cultured for 7 days maintained the low Bodipy^+^ phenotype and their phagocytic capacity using pHrodo *E.coli* bioparticles as before. Microglia with Bodipy^+^ lipid droplets and scarce phagocytosed bioparticles were hardly observed in microglia cultures obtained from repopulated old mice compared with cultures obtained in parallel from untreated old mice (Fig. 5E). These experiments support the concept that lipid droplet enrichment reduces the cell’s capacity to accumulate phagocytosed materials, suggesting that microglia repopulation in old mice improves microglia function.

**Fig. 5.**
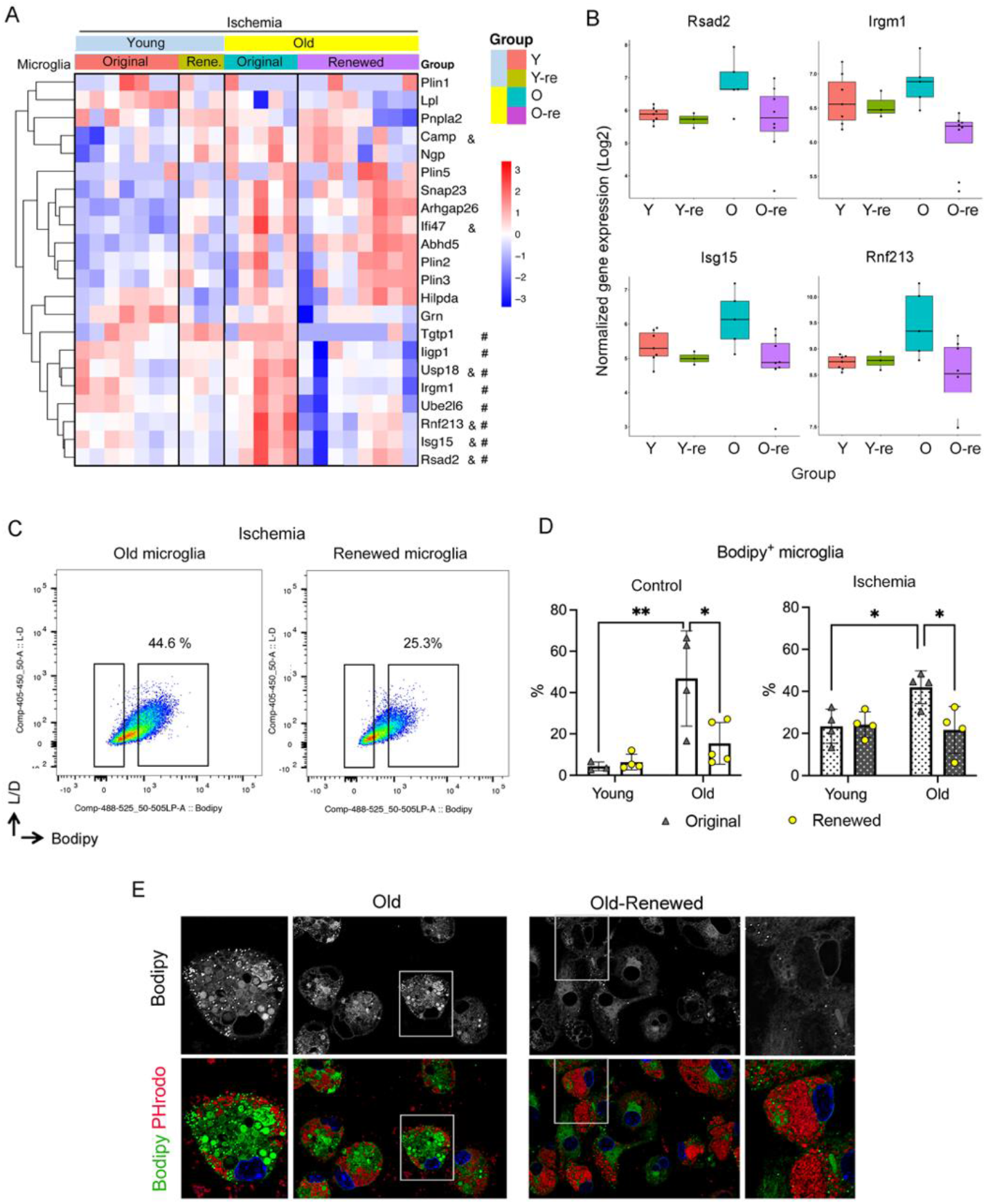
Microglia renewal eliminates the age-dependent subset of microglia containing lipid droplets. A) Heatmap of lipid droplet-associated proteins in microglia of young (n=7) and old (n=5) female mice and corresponding groups with microglia renewal (Re) under ischemia (n=3 and n=8). Old microglia show increased expression of some genes versus young microglia (&p<0.05). Microglia renewal in old mice prevents the age-dependent increase in type I IFN genes (#p<0.05). B) Box plots depict normalized gene expression of representative genes encoding lipid droplet-associated proteins of the IFN pathway. C) Representative flow cytometry plots showing Bodipy^+^ CD11b^+^CD45^low^ microglia from old male mice with or without microglia renewal (n=4 per group). Data show results from the ischemic and contralateral non-affected (control) brain hemispheres. In control, 40% of old microglia were Bodipy^+^, versus less than 5% for young microglia (**p=0.0053). Microglia renewal reduced the percentage of Bodipy^+^ microglia in old mice (*p=0.0172). After ischemia, 40% of microglia were Bodipy^+^ in old mice, whereas young ischemic mice showed 20% of Bodipy^+^ microglia (*p=0.0412), and microglia renewal reduced Bodipy^+^ microglia in old mice (*p=0.0242) (two-way Anova by age and condition and Tukey’s multiple comparisons test). Values show individual mice and bars show the mean±SD. Data for all the mice were obtained in parallel but for presentation purposed data for non-repopulated microglia are shown in Fig. 2D and Fig. 3F. D) Representative flow cytometry plots illustrating Bodipy^+^ cells within the CD45^low^ CD11b^+^ microglia population of ischemic tissue. E) Primary microglia cultures were obtained using CD11b^+^ magnetic beads from old male mice with or without microglia renewal (n=2 per group). Cells were exposed to red fluorescent pHrodo *E. coli* bioparticles and were stained with green fluorescent Bodipy and studied by confocal microscopy. Black and white images illustrate raw intensity data of Bodipy staining. The lower images are corresponding merged channels showing Bodipy in green, phagocytosed bioparticles in red, and the DAPI^+^ nuclei in blue. Repopulated microglia show less lipid droplets and the cytoplasm is replenished with phagocytosed bioparticles. The square regions of the images indicate areas magnified at the side of each image. Scale bar = 20 μm.

### Microglia renewal improves the neurological outcome after stroke

To investigate whether renewal of microglia in old mice affected the neurological dysfunction induced by stroke we conducted several behavioral tests in a longitudinal manner up to 14-days post-ischemia in mice treated with control diet or subjected to microglia depletion and repopulation (Fig. 6A). Compared with mice receiving control diet, microglia renewal ameliorated the neurological score and improved forelimb strength, as assessed by the grip test (Fig. 6B). The latency to fall in rotarod, which assesses motor activity, was higher (indicating better performance) in the repopulated group, but group differences were not statistically significant. The laterality index, as assessed with the cylinder test, showed a trend towards positive laterality, indicating impairment of the affected limb, in the control group at day four post-ischemia but not in the repopulated group (Fig. 6C). The volume of the brain lesion, as assessed by MRI at day four post-ischemia was similar in both groups (Fig. 6D), like the mean body weight before ischemia and one day later (Supp. Fig. S5A, B). At day two, mean body weight dropped more in the control group, but differences were not statistically significant (Supp. Fig S5A). Of note, we did not detect behavioral differences between diet groups during treatment prior to ischemia in old mice (Supp. Fig. S5C-D). Overall, old mice with renewed microglia showed signs of improvement in the neurological function after stroke.

**Fig. 6.**
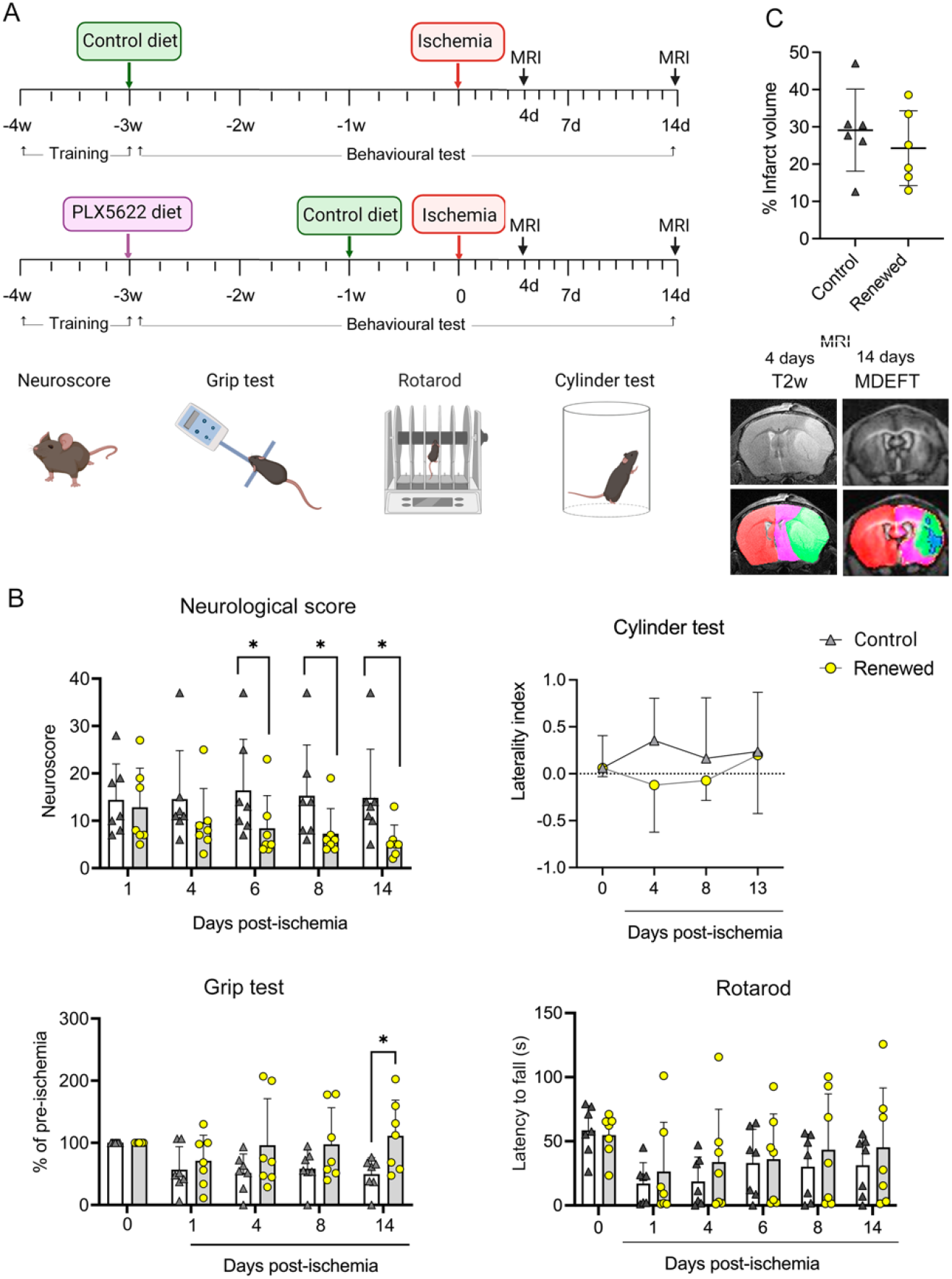
Microglia repopulation ameliorates the neurological outcome of stroke. A) Experimental design for microglia depletion and repopulation in old (21-22 month) female mice. We depleted microglia by providing a diet containing the CSF1R antagonist PLX5622 for three weeks. Microglia repopulation was induced by switching to control diet for seven days prior to induction of ischemia. Treatment controls were fed a corresponding control diet (n=7 per group). Behavioral testing (including training sessions) was conducted throughout the experiment and mice underwent two MRI studies. Measures were obtained the day prior to ischemia (basal value, time=0) and at different time points post-ischemia until day 14. B) The neurological score was lower, i.e., it improved, in mice with repopulated microglia (Mann-Whitney test, *p=0.0431 at day 6, *p=0.0280 at day 8, *p=0.0338 at day 14). The grip test assessing limb strength was higher in the group with repopulated microglia (Two-way Anova and Šídák’s multiple comparisons test, *p=0.0418). Values are expressed as % of the corresponding basal value (pre-ischemia) for each mouse. Rotarod test values are expressed as latency to fall (seconds) from the running wheel at different time points post-ischemia (higher means better). Positive values in the laterality index, as assessed via the Cylinder test, indicate impairment of the affected hindlimb, i.e., contralateral to the injured brain hemisphere. A trend towards positive laterality was only observed in the control group at day 4 post-ischemia (p=0.062 control group; p=0.578 repopulated group; Wilcoxon Signed Rank Test). One animal of the control group died after day four and received the worst score. Bars represent mean ± SD. C). The MRI lesion volume, as illustrated in the corresponding false color images below, showed no differences in infarct volume in mice with or without microglia renewal. Results correspond to day four postischemia. Representative MRI images obtained on day four post-ischemia using T2w MRI and day 14 using a Modified Driven Equilibrium Fourier Transform (MDEFT) MRI sequence (7.0 T BioSpec 70/30, Bruker BioSpin). Values show data for individual mice and/or the mean±SD.

## Discussion

Worse stroke outcome in elderly people is attributed to increased vulnerability of the aging brain to ischemia, but knowledge of the underlying biology is still poor^1^. Here we show that dysfunctional microglia contribute to the worse outcome of ischemic stroke in old mice. The underlying mechanisms involve an exaggerated type I IFN signature associated with age-dependent lipid droplet accumulation in microglial cells. Renewing the microglia population of old mice increases the overall fitness of these cells by reducing the subset of microglia containing lipid droplets. After stroke, repopulated microglia in old mice display attenuated innate immune responses and reduced lipid droplets compared with the original microglia of old mice. Importantly, old mice with repopulated microglia show improvement in the neurological dysfunction caused by stroke.

Microglia derived from erythromyeloid progenitor cells of the yolk sac colonize the mouse brain at embryonic stages^36–38^. These cells remain in the adult brain with negligible replacement by peripheral cells, and a low rate of self-renewal under steady state. In humans, microglial cells can live for more than twenty years^39^. Overall, the microglia lifespan is long, and cells display specific phenotypes in the aged brain^6,40^, which display dysfunctional microglia^8,11,41,42^. A hallmark of aging is the accumulation of lipofuscin inclusions^43,44^, which are found in several cells including microglia^41,42^. These lipofuscin pigment deposits are autofluorescent intracellular bodies containing lipid residues derived from the oxidation of polyunsaturated fatty acids and proteins^45^. Although there are different kinds of lipofuscin depending on the cell type, tissue, and disease condition, they often correspond to still active lysosomes containing hydrolytic enzymes. A previous study found that microglia repopulation in old mice restores the age-dependent dysfunction in lysosomal structure and intracellular lipofuscin accumulation^24^. Importantly, the increased myelin degradation by microglia in the aged brain is a recognized trigger of lipofuscin-like lysosomal inclusions^42^. Lysosomes carry out lipid hydrolysis and recruit lipid droplets to generate free fatty acids. We observed lysosomes attached to lipid droplets in microglia after ischemia, and similar observations were reported in microglia of old rats^46^. These findings suggest cellular lipid recycling in microglia through lipophagy, i.e. a form of autophagy, both after brain ischemia and in aged subjects. Moreover, the transcriptomic profile of old versus young microglia after ischemia showed enrichment of metabolic pathways related to lipid homeostasis, whereas pathways involved in peroxisomal beta-oxidation of very long chain fatty acids were downregulated. Altogether, these findings are compatible with age-dependent impairment of the capacity to shorten the chain of very long chain fatty acids derived from lysosomal lipid degradation^47^, thus suggesting alterations in lipid disposal in microglia of aged subjects. In turn, we found that repopulated microglia in old mice after stroke show transcriptional enrichment of pathways related to lipid metabolism and have less lipid droplet accumulation. Altogether, the current findings in repopulated microglia demonstrate ‘rejuvenation’ of certain metabolic features typically associated with microglia of old mice.

Lipid droplets originate from the endoplasmic reticulum and store neutral lipids that can be later metabolized via β-oxidation to generate ATP. After bacterial infection, hepatocytes generate lipid droplets surrounded by anti-microbial proteins^29^. Lipid droplets are critically involved in mounting an efficient IFN response against different types of infection^29, 30, 48–50^. Our results show lipid droplet biogenesis associated with the innate immune response induced by brain ischemia in microglia. Of note, aging and ischemia upregulated Viperin, which can promote TLR7- and TLR9-mediated production of type I IFN in plasmacytoid dendritic cells^51^. Microglial cells rich in lipid droplets showed less phagocytosed *E. coli* bioparticles supporting the reported impaired phagocytic function of microglia containing lipid droplets^11^. The accumulation of lipid droplets seems to contribute to microglia dysfunction, impairing the cellular fitness for tissue clearance, in contrast to the plausible benefits of lipid droplet accumulation in other cells after infection^29,48^.

This study also demonstrates that stroke induces lipid droplet biogenesis in microglia of young mice within hours/days after stroke onset. Therefore, given that accumulation of lipid droplets is a phenotypic trait of old microglia, stroke causes accelerated and premature ‘aging’ of microglia in just a few days^11^. Very recent studies reported ischemia-induced changes in lipid metabolism in microglia,^52^ and accumulation of lipid droplets in chronic stroke lesions impairing stroke recovery^53^. Our results show that microglia accumulate lipid droplets shortly after stroke onset due to lipid enrichment plausibly attributable to the innate immune response, phagocytosis of damaged tissue, and metabolic disturbances that are associated with the increased expression of certain lipid-droplet associated proteins, such as Plin2. Brain ischemia causes tissue necrosis and inflammation, increasing the requirement of phagocytosis to remove dead cells, tissue debris, myelin fragments, and infiltrating leukocytes^18^. During this process, we show that microglia develop a strong type I IFN signaling response likely attributable to recognition of intracellular danger signals, e.g., via the cGAS/Sting pathway^54,55^, thus upregulating the expression of innate immune molecules associated to lipid droplets. The post-ischemic IFN response is further exacerbated in microglia of old versus young mice, whereas it is reduced in repopulated microglia of old mice to the levels of microglia of young mice, in consonance with the reduction in age-associated lipid droplets.

We do not know whether all microglial cells are able to generate lipid droplets under specific conditions; however, we observed that microglia devoid of lipid droplets in the aged mice did not generate new lipid droplets after ischemia, unlike Bodipy^neg^ microglia of young mice, suggesting that there might be subsets of microglia prone or resistant to lipid droplet generation. Microglia heterogeneity has been described depending on age, brain region, and injury/disease^6,56–58^. Therefore, it is possible that the regional location contributed to the abundance of Bodipy^+^ microglia since, for instance, aging-related lipofuscin accumulation associated with myelin degradation is more prominent in the white matter than in the grey matter ^42^.

Strategies designed to remove age-related dysfunctional microglia in the elderly could ameliorate microglia function, increasing the cellular resilience to cope with brain injury induced by stroke and possibly other neurological conditions. Concerns about putative safety issues related to the process of microglia renewal demand further investigation.

## Acknowledgments

Study funded by the Spanish Ministry of Science and Innovation (MICINN), co-financed by Fondo Europeo de Desarrollo Regional (FEDER) (grants SAF2017-87459-R and PID2020-113202RB-I00). PTI-Neuro-Aging Platform of the Spanish National Research Council (CSIC) supported this work and funded MG. MAR has a predoctoral fellowship MICINN-FPI (PRE2018-085737). We thank the technical support of Dr. Eva Prats and Josep Rebled of the Electron Microscopy Service-*Campus Facultat de Medicin*a, *Serveis Cientifico-Tècnics de la Universitat de Barcelona*. We acknowledge the help of Dr. Valérie Petegnief and the technical support of Leonardo Márquez-Kisinousky and Francisca Ruiz-Jaén. We acknowledge the support of the Cytomics and MRI imaging facilities of IDIBAPS. RNA sequencing was performed at the *Centro Nacional de Análisis Genómico* (CNAG) in the Centre of Genomic Regulation (CRG) of Barcelona. Part of the work was performed at Centre de Recerca Biomèdica Cellex. The *Centres de Recerca de Catalunya* (CERCA) Program of Generalitat de Catalunya supports IDIBAPS. Plexxikon provided PLX5622 under Materials Transfer Agreement.

## Author Contributions

Conceptualization, A.M.P.; Methodology, M.G., M.A.R., C.C.; Investigation, M.A.R., M.G., and J.P.; Formal Analysis: J.L.; Data Curation, J.L; Writing – Original Draft, A.M.P.; Writing - Review & Editing, M.G., M.A.R., J.P., A.P., Visualization, M.A.R., M.G., A.M.P.; Supervision, A.M.P., M.G., A.P.; Funding acquisition, A.M.P.

## METHODS

### Materials & Correspondence

Further information and requests for resources and reagents should be directed to and will be fulfilled by the corresponding author, Anna M.Planas.

### Mice

We used adult young (3-4 month) mice and old (21-22 month) mice in the C57BL/6 background obtained from Janvier (Lyon, France) or Envigo (Amsterdam, The Netherlands). We also used CX3CR1creERT2 mice (B6.129P2(C)-Cx3cr1tm2.1(cre/ERT2)Jung/J, #SN020940) crossed with reporter ROSA26-tdTomato mice (B6.Cg-Gt(ROSA)26Sortm9(CAG-tdTomato)Hze/J, #SN007909) obtained from The Jackson Laboratory. The latter mice received tamoxifen three weeks before surgery. For generating chimeric mice, we used donor DsRed-fluorescent reporter mice from our colony maintained at the animal house of the School of Medicine of the University of Barcelona (UB). We used female and male mice, but we did not mixed sexes in the same experiments. For clarity, the sex of the mice is reported in the figure legends. Animal work was conducted following the Catalan and Spanish laws (Real Decreto 53/2013) and the European Directives. All experiments were conducted with approval of the ethical committee (Comité Ètic d’Experimentació Animal, CEEA) of UB, and the local regulatory bodies of the Generalitat de Catalunya, and in compliance with the NIH Guide for the Care and Use of Laboratory Animals. The work is reported following the ARRIVE guidelines.

### Induction of brain ischemia

Cerebral ischemia was induced by transient occlusion of the right middle cerebral artery (MCAo) with the intraluminal technique, as described^26^, with modifications. In brief, anesthesia was induced with 4% isoflurane in a mixture of 30% O_2_ and 70% N2O and it was maintained with 1%–1.5% isoflurane in the same mixture using a facial mask. With the animal in supine position on a heated surgery table (37°C), a longitudinal cut was produced in the ventral midline of the neck and the submaxillary glands and the omohyoid and sternothyroid muscles were separated, exposing the right carotid artery. A monofilament (#701912PK5Re, Doccol Corporation, Sharon, MA) was introduced through the right external carotid artery up to the level where the MCA branches out. We allowed the mice to wake up and kept them on a heated blanket at 37°C. After 45 min of arterial occlusion, mice were anesthetized again, the filament was cautiously removed, and the suture of the ipsilateral CCA was removed to allow reperfusion. CBF was monitored with laser Doppler flowmetry (Perimed, AB, Järfälla, Sweden) during the procedure to guide the filament insertion and monitor perfusion. Mice received analgesia (buprenorphine, 150 μl, 0.015 mg/mL, via s.c.) and were kept on a thermal blanket at 37°C for 1 hour after surgery. Pre-established exclusion criteria were either unsuccessful MCAo resulting in no infarction or technical surgical complications. MCAo was carried out under isoflurane anesthesia using a monofilament (#701912PK5Re, Doccol Corporation, Sharon, MA).

### Drug treatments

For microglia depletion, mice received the CSF1R inhibitor PLX5622 (Plexxikon Inc, Berkeley USA) following previously reported protocol^18^. PLX5622 was mixed into AIN-76A rodent diet at 1,200 ppm (Brogaarden, Denmark). Treatment controls received AIN-76A drug-free standard chow for the same period. Both diets were given in parallel in groups of 5 animals per cage. Mice received the diet ad libitum. In as much as possible, we blinded the treatment, e.g. for evaluation of behavioral data. The various dosing regimens are explained in the figures.

### Generation of chimeric mice

We generated chimeric mice by chemical ablation of the bone marrow of wild type (WT) recipient mice followed by transplantation of bone marrow from DsRed reporter donor mice, as described (Kierdorf et al., 2013). In brief, adult (2-month-old) male WT mice received three intraperitoneal injections of the chemotherapeutic agent busulfan (30 μg/g body weight) 7, 5 and 3 days prior to the transfer of five million bone marrow cells from DsRed donor mice via the tail vein. The mice were studied at least 8 weeks after transplantation.

### Assessment of Neurological function

We used a composite neuroscore modified from a previously reported neuroscore^60^. The score ranged from 0 (no deficits) to 37 (poorest performance in all items) and it was calculated as the sum of general and focal deficits, including the following general deficits (scores): hair (0– 2), ears (0–2), eyes (0–3), posture (0–3), spontaneous activity (0–3), and epileptic behavior (0– 1); and the following focal deficits: body symmetry (0–2), gait (0–4), climbing (0–3), circling behavior (0–3), forelimb symmetry (0–4), compulsory circling (0–3), and whisker response (0– 4). For behavioral tests, mice received preoperative training prior to starting drug treatments, and were tested during treatments to check for possible behavioral alterations. Tests were performed at different time points prior and after ischemia, and the last test performed prior to ischemia was taken as the basal value. The rotarod test (Rota-Rod/RS Panlab Harvard apparatus) assesses motor activity. We carried out three trials on an accelerating rod, starting at 4 rpm with an increasing acceleration of 1 rpm each 8 s with 15 min of rest between the trials. The latency before failing was taken as the average of the three trials. The grip test measures maximal muscle strength of forelimbs when the mouse loses its grip on the T-bar. A computerized grip strength meter (Alemo 2450, Ahlborn) was used. We performed three measures each testing day and we used the best measure. The laterality index was calculated with the cylinder test. In brief, the mouse was introduced in a plexiglass cylinder with an internal diameter of 9.5 cm and a height of 15 cm, and the mouse was videorecorded for 10 min. We counted in the videos the number of times that the mouse raised the body and touched the cylinder (first touch only) for vertical exploration of the cylinder. We counted the number of first contacts with the right (non-affected) or left (affected) forelimb, or both simultaneously. The laterality index was calculated as the number of contacts with the right limb minus those with the left limb, divided by the total number of first touches. Two independent observers analyzed the videos and the mean value was calculated.

### Magnetic Resonance Imaging (MRI)

The brain lesion was imaged using MRI in a 7.0 T BioSpec 70/30 horizontal animal scanner equipped with a 12-cm inner diameter actively shielded gradient system (400 mT/m). The receiver coil was a phased array surface coil for mouse brain (Bruker BioSpin, Ettlingen, Germany). Four days and 14 days after induction of ischemia, the brain was studied with T2w turbo RARE fast spin-echo MRI sequences with 1 effective echo time (ET)= 33 ms, slice thickness = 0.5 mm, repetition time (TR) = 2336 ms, field of view = 20 × 20 mm^2^, matrix size = 256 pixels, and in-plane spatial resolution = 0.078 mm. For evaluation of structural alterations that were no longer observable in T2 relaxometry maps at day 14 post-ischemia, we acquired high resolution 3D Modified Driven Equilibrium Fourier Transform (MDEFT) images. The scan parameters were as follows: TE = 2.2 ms, slice thickness = 0.5 mm, TR = 4000 ms, 4 segments, in-plane spatial resolution 0.078 mm. Images were obtained with ParaVision 6.0 software (Bruker BioSpin, Ettlingen, Germany). Image analysis was carried out using ImageJ.

### Brain tissue processing and Flow cytometry

Mice were anesthetized with isoflurane and euthanized by cervical dislocation. The brain was carefully collected, in order not to damage the tissue, and immersed in Hanks’ Balanced Salt solution w/o ions (HBSS w/o Ca^2+^ and Mg^2+^; #14175-053, Thermo Fisher Scientific) on ice. The forebrain was dissected with a scalpel, discarding the cerebellum and the olfactory bulbs, and minced in small pieces. The Neural Tissue Dissociation Kit - P (NTDK, #130-092-628, Miltenyi Biotec) based on the enzymatic dissociation with papain was used to homogenize the tissue. Mechanical dissociation was performed with the gentleMACS™ Octo Dissociator (#130-096-427, Miltenyi Biotec: 1x m_Brain_1 program and 1x ABDK_37C program by Miltenyi) with the tissue immersed in the buffers and enzymes from the NTDK kit, according to manufacturer instructions. The tissue was then filtered through a 70 μm cell strainer (#352350, Falcon) previously humidified with HBSS with Ca^2+^ and Mg^2+^ (#14025-092, Thermo Fisher Scientific). Then, cells were separated from myelin by an immunomagnetic separation method. Brain cells were incubated with Myelin Removal Beads II (#130-096-733, Miltenyi Biotec) and then passed through LS Columns (#130-042-401, Miltenyi Biotec) held onto the OctoMACS Separator (#130-091-051, Miltenyi Biotec) and to the MACS^®^ MultiStand (#130-042-303, Miltenyi Biotec). Unspecific binding of antibodies was blocked by previous incubation for 10 min with anti CD16/CD32 (Fc block, clone 2.4G2; #55314, BD Pharmingen) in Fluorescence Activated Cell Sorting (FACS) buffer at 4 °C. Live/dead Aqua cell stain (#L34957, Thermo Fisher Scientific) was used to determine the viability of cells. Cells were incubated during 30 minutes at 4°C with the following primary antibodies: CD11b (clone M1/70, AF647, #557686, BD Pharmingen), CD45 (clone 30-F11, Brilliant Violet 786, #564225, BD Horizon). After washing once with phosphate buffered saline (PBS), cells were stained with BODIPY (4,4-difluoro-3a,4adiaza-s-indacene, Molecular Probes BODIPY 493/503, #D3922, Life Technologies) diluted 1:1,000 during 15min at RT. After a wash with cold FACS buffer, the cells were analyzed in a BD FACSAriaII cytometer using FacsDiva software (version 5, BD Biosciences, San Jose, CA, USA). Data analyses were performed with FlowJo software (version 10, FlowJo LLC, Ashland, OR, USA).

### Cell isolation

Microglia for the ischemic vs. control (data in Fig. 1) was isolated from the brain of CX3CR1cre^ERT2^-ROSA26 tdTomato (tdT) mice via FACS after intracardiac perfusion with 2ml of cold PBS. Brains were collected in cold HBSS buffer (w/o Ca^2+^ and Mg^2+^; #14175-053, Thermo Fisher Scientific). The brain tissue was enzymatically dissociated using the Neural Tissue Dissociation Kit (P) (#130-092-628, Miltenyi Biotec). The gentleMACS™ Dissociator (#130-096-427, Miltenyi Biotec) was used for mechanical dissociation steps following the Neural Tissue Dissociation Kit (P) manufacturer protocol for dissociation without heaters. The digested tissue was filtered twice with 70 μm and 40 μm filters, washing with cold Hanks’ balanced salt solution (HBSS) (with Ca^2+^ and Mg^2+^; #14025-092, Thermo Fisher Scientific). Cells were separated from myelin and debris by 30% isotonic percoll gradient (#17-0891-01, GE Healthcare) in Myelin Gradient Buffer (MGB). Samples were centrifuged at 950 *xg* without acceleration or brake, for 30 min at 4°C. Cells were collected from the bottom of the tube, washed once with cold FACS Stain Buffer (#554656, BD Biosciences), and processed for FACS in a FACSAriaII sorter (BD Biosciences). No staining was required since the expression of tdT was used to separate microglial cells.

Microglia for the old vs. young comparisons was isolated from the brain of WT mice after the assigned depletion/renewal protocol and ischemia. Mice were euthanized under deep anesthesia and the brain was processed in the same way as for the flow cytometry experiments to obtain a single cell suspension (see above). Unspecific binding was blocked by incubation for 10 min with anti CD16/CD32 (in FACS buffer at 4°C. Live/dead Aqua cell stain was used to determine the viability of cells. Cells were incubated with the following primary antibodies during 30 minutes at 4°C: CD11b (clone M1/70, APC-Cy7, #557657, BD Pharmingen), CD45 (clone 30-F11, FITC, #553080, BD Pharmingen). After washing with FACS Stain Buffer (#554656, BD Biosciences), the cells were sorted in a FACSAriaII SORP sorter (BD Biosciences). Microglial cells were collected in sterile DPBS (#14190-094, Thermo Fisher Scientific), centrifuged, and resuspended in lysis buffer (from PureLink™ RNA Micro Kit #12183016, Invitrogen) supplemented with 10% β-mercaptoethanol and finally snap-frozen in dry ice.

### Adult microglia culture

Microglia cells from adult young or old mice were isolated and cultured using immunomagnetic separation (Miltenyi Biotec, Germany). Mice were perfused via the left ventricle with 60 mL of cold saline and collected in HBSS buffer without calcium/magnesium (#14175-05; Life Technologies). The brain tissue was enzymatically dissociated using the Neural Tissue Dissociation Kit-P (as above). The gentleMACS™ Dissociator with Heaters (#130-096-427; Miltenyi Biotec) was used for mechanical dissociation steps during 30 minutes at 37ºC. The digested tissue was filtered (70μm) with HBSS buffer with calcium and magnesium and prepared for myelin removal process (Myelin Removal Beads II, #130-096-733; Miltenyi Biotec). Then, cells were magnetically labeled with CD11b microbeads (#130-093-634; Miltenyi Biotec) diluted in PBS supplemented with 0.5% BSA for 15 minutes in the dark in the refrigerator (2-8 °C). CD11b^+^ cells were collected using magnetic field columns (Miltenyi Biotec). Cell suspensions (35 μl) were then plated in complete medium consisting of DMEM medium (#10569010; Gibco-BRL) supplemented with 10% fetal bovine serum (FBS; Gibco-BRL) and containing 40U/mL penicillin and 40μg/mL streptomycin (#15140122; Gibco-BRL) added as a drop in the middle of each well of a Poly-L-Lysine (#P4832; Sigma) pre-coated 8-well plate (μ-Slide 8 Well, IBIDI #80826). Cells were incubated for 30 minutes at 37 °C and then 250 μL of complete medium were carefully added to each well. 24 hours later, we replaced 50% of complete medium, and we did a full medium change at day 5. The cells were maintained at 37 °C in a humidified atmosphere of 5% CO_2_ for 7 DIV.

To study phagocytosis we used red fluorescent pHrodo *E.coli* bioparticles (#P35361, Thermo Fisher Scientific) in the phagocytosis assay. At 7 DIV, microglial cells were exposed to the fluorescent beads (0.1 mg of particles/mL) for 1h or 4h. Following 3 washes with PBS pH 7.4 to remove all the non-phagocytosed particles, cells were fixed with cold 4% paraformaldehyde for 10 min. After two washes with PBS, fluorescent Bodipy (Molecular Probes BODIPY 493/503, #D3922, Life Technologies) was added (1:1,000) for 10 minutes at room temperature and protected from the light. DAPI (#D3571, Life Technologies) stain was performed to visualize the cell nuclei. Two more washes with PBS were completed before proceeding to image visualization. Images were obtained with a confocal microscope (Dragonfly, Andor) and were not further processed if not to enhance global signal intensity in the entire images of the same experiment for image presentation purposes using ImageJ or Adobe Photoshop.

### Immunofluorescence in brain tissue sections

Mice were perfused through the heart with 40 mL of cold saline (0.9%) followed by 20 mL of cold 4% paraformaldehyde diluted in phosphate buffer (PB) pH 7.4. The brain was removed, fixed overnight with the same fixative, and immersed in 30% sucrose in PB for cryoprotection for at least 48 h until the brains were completely sunk to the bottom of the falcon. After that, brains were frozen in isopentane at −40ºC. Cryostat brain sections (14-μm thick) were fixed in ethanol 70%, blocked with 3% normal serum, and incubated overnight at 4ºC with the primary antibody: rabbit polyclonal antibody against P2RY12 (1:250, #AS-55043A, AnaSpec Inc.). The secondary antibody was: Alexa Fluor 488 (Molecular Probes; Life Technologies S.A.). Cell nuclei were stained with DAPI or To-Pro3 (Invitrogen). Confocal images were obtained (TCS-SPE-II microscope, Leica Microsystems).

### Transmission electron microscopy

Mice were perfused through the heart with 20 mL of phosphate buffer 0.1M pH 7.4 followed by 20 mL of the fixative solution (mixture of paraformaldehyde 2% and glutaraldehyde 2.5% in phosphate buffer 0.1M pH 7.4). Small pieces of 1,5 mm3 of cortex from the core and penumbra, and of striatum were obtained and post-fixed in osmium tetroxide (1%) and potassium ferrocyanide (0.8%), dehydrated with acetone and embedded in Spurr resin. Ultrathin sections (60-80 nm) were obtained with an ultramicrotome (Leica Ultracut E) using a diamond knife (Diatome). Sections were mounted on copper grids, stained with 2% uranyl acetate for 10 min and lead citrate 2 min, and observed in a Transmission electron microscope JEOL 1010 with an Orius (Gatan) CCD Camera.

### Western blotting

Mice were euthanized under isoflurane anesthesia brains were carefully removed from the skull and perfused through the heart as above. The ipsilateral and contralateral brain hemispheres were dissected out, immediately frozen on dry ice, and stored at −80 °C until further use. Tissue was homogenized in radioimmunoassay precipitation (RIPA) buffer, centrifuged for 20 min at 12,000 *xg* at 4 °C and the supernatant was used for protein determination by the Bradford method. Twenty-five μg of protein were mixed with loading buffer containing β-mercaptoethanol and samples were loaded in 10% polyacrylamide gels for electrophoresis. Proteins were transferred to polyvinyl difluoride membranes (Immobilon-P, #IPVH00010, Millipore/Sigma) and incubated overnight at 4 °C with the primary anti-Plin2 antibody (ADFP rabbit polyclonal antibody, # PA1-16972, Invitrogen) diluted 1:500, followed by horseradish peroxidase-conjugated secondary antibody 1h at RT. β-Tubulin (#T4026, Sigma-Aldrich; 1:100,000) was used as loading control. Blots were developed with a chemiluminiscent substrate (ECL Amersham, # RPN2235).

### RNA Extraction

RNA was extracted from samples of FACS-sorted microglia with PicoPure™ RNA Isolation Kit (#KIT0204, Thermo Fisher Scientific) following manufacturer instructions with minor modifications. RNA was precipitated with 70% ethanol. To avoid genomic DNA contamination a DNAse step was performed using Invitrogen™ PureLink™ DNase Set (#12185010, Invitrogen). RNA quantity and purity were assessed with the Pico Kit Assay on the Agilent 2100 Bioanalyzer System.

### RNA sequencing

Libraries were prepared using NEBNext^®^ Poly(A) mRNA Magnetic Isolation Module (# e7490) and NEBNext^®^ Ultra II Directional RNA Library Prep Kit for Illumina (24 reactions # e7760 or 96 reactions # e7765) according to the manufacturer’s protocol, to convert total RNA into a library of template molecules of known strand origin and suitable for subsequent cluster generation and DNA sequencing. Briefly, 10 ng to 50ng of total RNA were used for poly(A)-mRNA selection using poly-T oligo attached magnetic beads using two rounds of purification. During the second elution of the poly-A RNA, the RNA was fragmented under elevated temperature and random primed. Then, the cleaved RNA fragments were copied into first strand cDNA using reverse transcriptase. After that, second strand cDNA was synthesized, removing the RNA template and synthesizing a replacement strand, incorporating dUTP in place of dTTP to generate ds cDNA using DNA Polymerase I and RNase H. These cDNA fragments, then were ‘A’-tailed, NEB hairpin adaptor was ligated and USER enzyme was used to excise the loop on the hairpin adaptors. Finally, PCR selectively enriched those DNA fragments that had adapter molecules on both ends and to amplify the amount of DNA in the library and to add specific barcodes to each sample. Final libraries were analyzed using Bioanalyzer DNA 1000 or Fragment Analyzer Standard Sensitivity (# 5067-1504 or # DNF-473, Agilent) to estimate the quantity and validate the size distribution and were then quantified by qPCR using the KAPA Library Quantification Kit KK4835 (# 07960204001, Roche) prior to the amplification with Illumina’s cBot. Libraries were sequenced 1 * 50+8 bp on Illumina’s HiSeq2500.

### Transcriptomic analysis

Analyses were performed as previously reported^61^. In brief, raw reads were analyzed for data quality using FastQC v0.11.5 (Babraham Bioinformatics) and filtered using skewer v0.2.2^62^ for removing the low-quality reads and trimming the Illumina adapter. STAR program^63^ against Mus musculus genome (GRCm38) was used for mapping the reads followed by the quantification of genes with the RSEM program^64^ using GENCODE m15 reference annotation^65^. After eliminating genes without an expected value greater than ten and non-autosomal, we used TMM method and limma-voom transformation^66^ to normalize the non-biological variability. Differential expression between different groups was assessed using moderated t-statistics^67^. Gene Ontology and Reactome canonical pathway enrichment analysis was performed through GSEA function in cluster Profiler package (gseGO and gsepathway for enrichment analysis and cnetplot graphics) using previously computed t-statistic. Heatmaps and Principal component plots were performed using R statistical software.

### Statistical analyses

Two-group comparisons were carried out with two-tailed Mann–Whitney test or t-test, as required after testing for normality, and adjusting for paired measures using Wilcoxon-matched-pairs signed rank test or paired t-test. Multiple groups were compared with one-way ANOVA or Kruskal-Wallis test, or two-way ANOVA, followed by appropriate post-hoc analyses. In experiments designed to test whether microglia repopulation in old mice could improve the neurological deficits induced by stroke, sample size was calculated using G*power 3.1 software (University of Düsseldorf) with an alpha level of 0.05 and a statistical power of 0.9. The size effect (d=2.08) was estimated based on differences in the neuroscore mean and SD between old and young mice after induction of ischemia. In other experiments, we made estimations based on information on the group mean and SD from previous flow cytometry or RNA data of our own laboratory, and we built from these data the number of animals needed for comparing other outcome measures. For measurements designed as proof of concept, validation, or as internal controls we used minimum reasonable numbers of animals for confirmatory purposes. The specific tests used in each experiment, p values, and n values are stated in Figure Legends. We used GraphPad Prism software version 9.3.1.

### Data availability

The RNA-Seq data are accessible from the GEO repository of the National Center for Biotechnology Information, U.S. National Library of Medicine (The accession numbers for these data are GEO: GSE136856 and GSE196737) (https://www.ncbi.nlm.nih.gov/geo/info/linking.html). Other datasets will be found in the repository of our Institution, CSIC, or will be available from the corresponding author upon reasonable request.

