## Supplemental figures for "Age-dependent lipid droplet-rich microglia worsen stroke outcome in old mice"

Supplementary Fig. S1

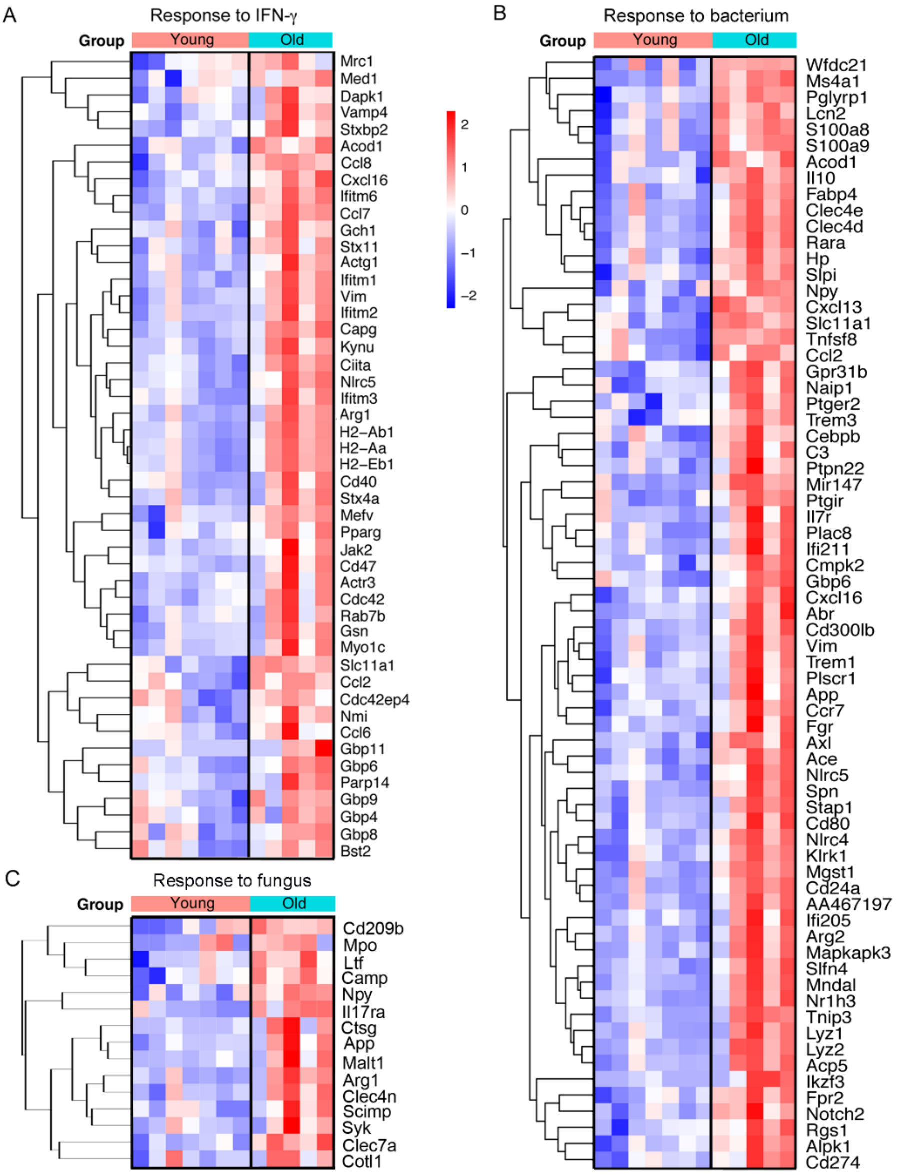


**Suppl. Fig. S1. Enrichment of genes in GO pathways involved in immune responses in microglia of old vs. young ischemic mice.** *Related to Fig. 3.* RNAseq analysis of microglia obtained by FACS from the brain of old (n=5) versus young mice (n=7) four days after ischemia. Heatmaps illustrate genes upregulated in microglia of old ischemic mice vs. young ischemic mice for the following GO terms: 'Response to IFN-γ' (A), 'Response to bacterium' (B), and 'Response to Fungus' (C).

Supplementary Fig. S2

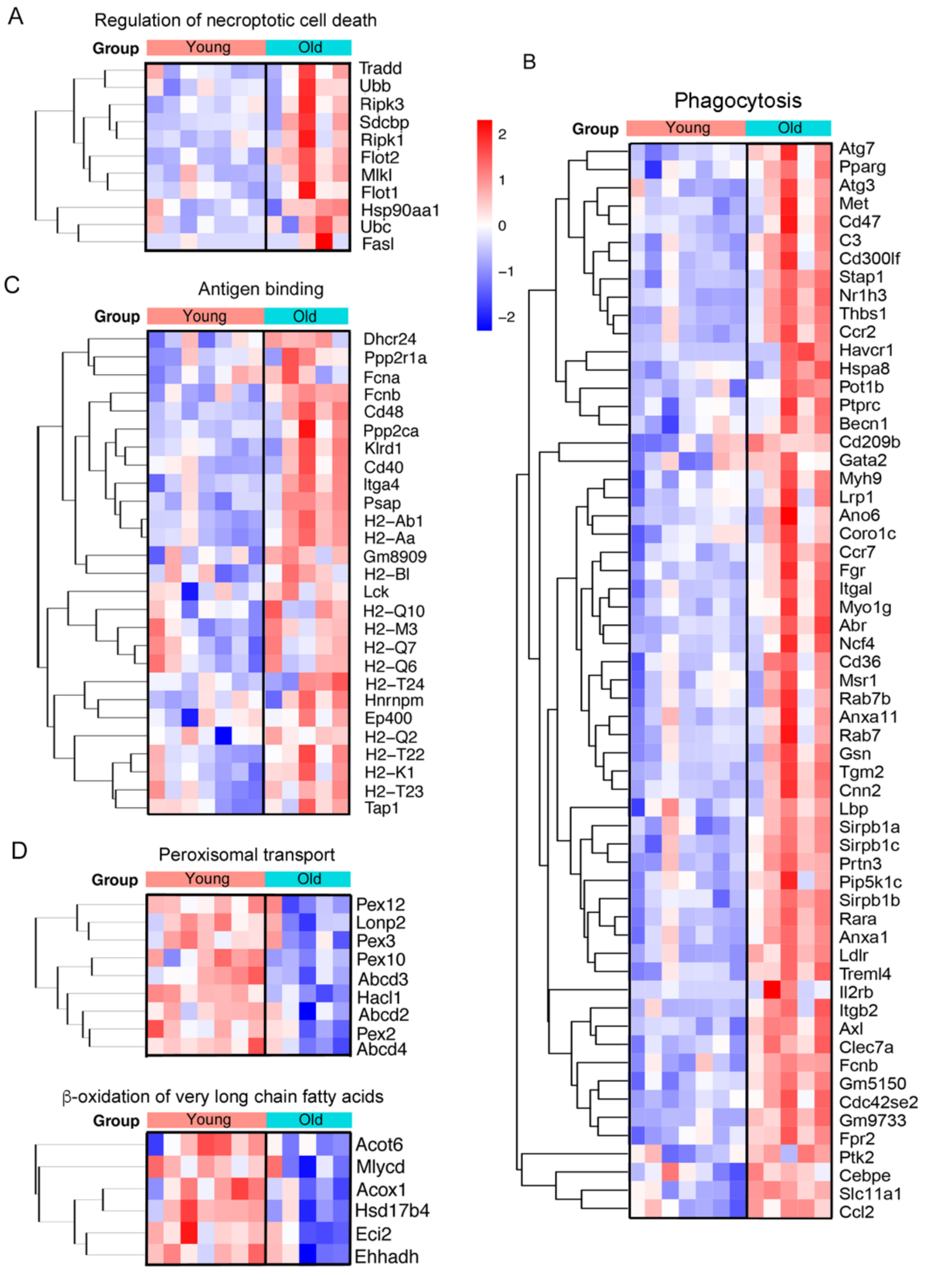


**Suppl. Fig. S2. Enrichment of genes in GO pathways involved in functional and metabolic responses in microglia of old vs. young ischemic mice.** *Related to Fig. 3.* RNAseq analysis of microglia obtained by FACS from the brain of old (n=5) versus young mice (n=7) four days after ischemia. Heatmaps illustrate genes upregulated in microglia of old vs. young mice after ischemia for the GO terms: A) 'Regulation of necroptotic cell death', B) 'Phagocytosis', and C) 'Antigen binding'. In contrast, downregulated pathways in microglia of old mice included: D) 'Peroxisomal transport' and 'β-oxidation of very long chain fatty acids'.

Supplementary Fig. S3

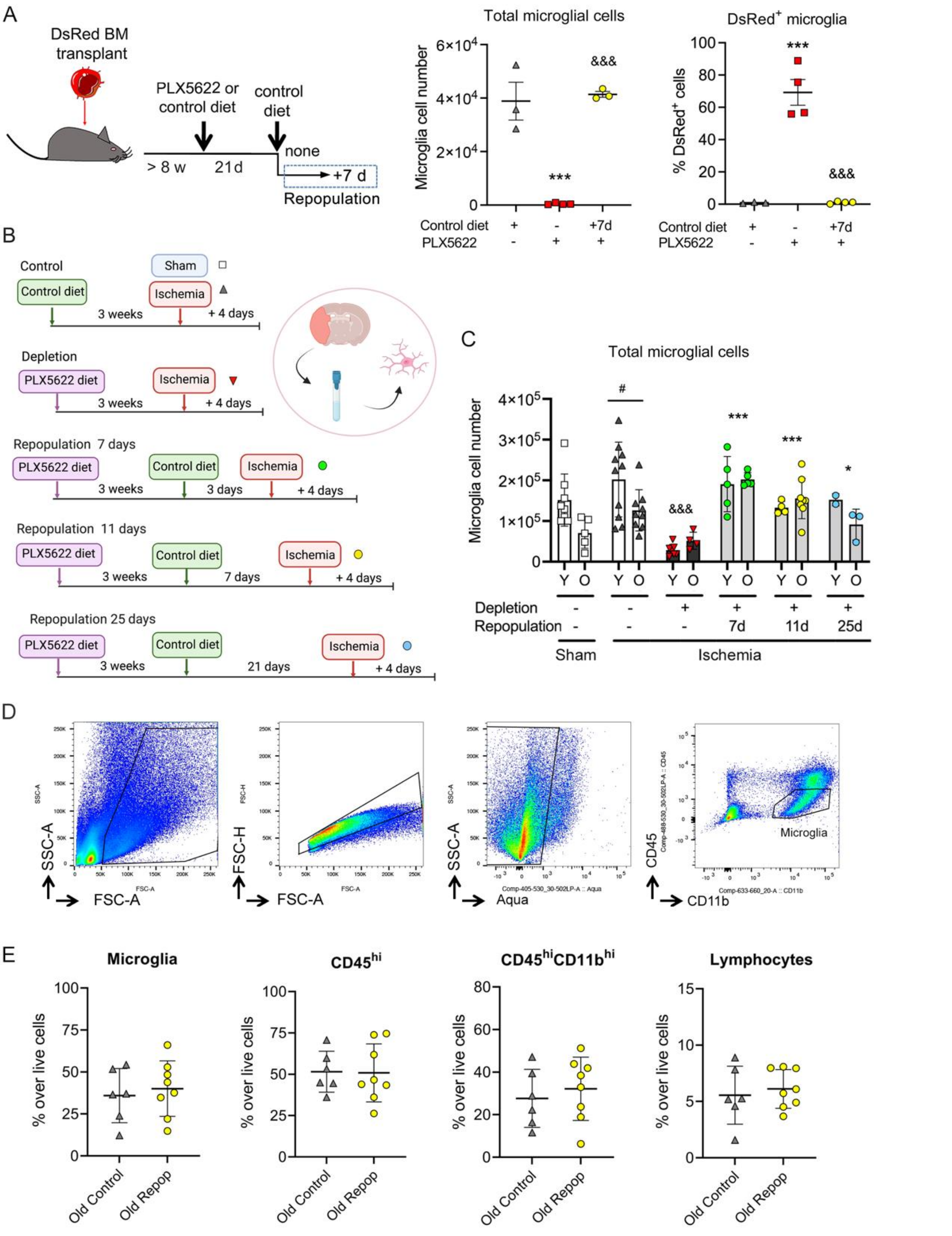


**Suppl. Fig. S3. Microglia cell number after depletion/repopulation.** *Related to Fig.4.* A) Renewed microglia derive from brain cells. We generated chimeric mice by bone marrow transplantation from DsRed fluorescent reporter donor male mice to wild type recipient male mice (n=11). After at least eight weeks, mice were treated with PLX5622 in the diet (n=8) or corresponding control diet (n=3). Three weeks later, mice were euthanized (n=4) or they were switched to control diet for repopulation for seven days (n=4 per group). The brain was studied via flow cytometry by measuring DsRed$^+$ and DsRed$^-$ cells in the gate of microglia cells. Absolute number of microglial cells was strongly reduced after microglia depletion (PLX5622 diet) (one-way

Anova and Holm-Šídák's multiple comparisons test ***p=0.0002 vs. control diet). Microglia numbers recovered after seven days of repopulation ($^{\&\&\&}$p=0.0002 vs. depleted cells). The % of DsRed cells in the microglia gate of mice fed control diet was negligible. However, after microglia depletion (PLX5622 treatment) (Anova and Holm-Šídák's, ***p<0.0001 vs. control diet) there was a high % of DsRed$^{+}$ cells within the very small population of CD45$^{low}$CD11b$^{+}$ cells indicating the presence of a few CSF1R-independent infiltrating cells. Importantly, the proportion of DsRed$^{+}$ cells was negligible after mice were switched to control diet and the number of microglia increased ($^{\&\&\&}$ p<0.0001 vs. depleted mice). B) Experimental design for microglia depletion and repopulation in young (3-4 month) (n=39) and old (20-22 month) (n=34) female mice. We depleted microglia via a PLX5622 diet for three weeks. Mice were repopulated by switching to the corresponding control diet for 3, 7, or 21 days prior to ischemia induction. The brain was studied four days post-ischemia and during this period control diet was maintained. Total repopulation times: 7, 11 and 25 days, respectively. As controls, we used mice subjected to control diet and studied four days post-ischemia or sham-operation. C) The number of microglia recovered post-ischemia was lower in old than in young mice fed a control diet (two-way ANOVA and Šídák's multiple comparisons test, *p=0.0134). The PLX5622 diet strongly reduced the number of microglial cells post-ischemia in both age groups ($^{\&\&\&}$ p<0.0001 vs. ischemic mice on control diet); switching to control diet increased the number of microglia vs. depleted mice (***p<0.0001 at day 7, ***p=0.0003 at day 11, and p=0.0377 at day 25) (two-way ANOVA and Dunnett's multiple comparison test). D) Gating strategy for cell sorting to obtain the CD45$^{low}$CD11b$^{+}$ microglia shown in (C). E) Microglia and brain infiltrating cells four days post-ischemia in old female mice under control diet (n=6) or following microglia depletion/repopulation for seven days prior to ischemia (n=8). There were no differences between groups regarding the % of cells. Values show data for individual mice and the mean ± SD.

Supplementary Fig. S4

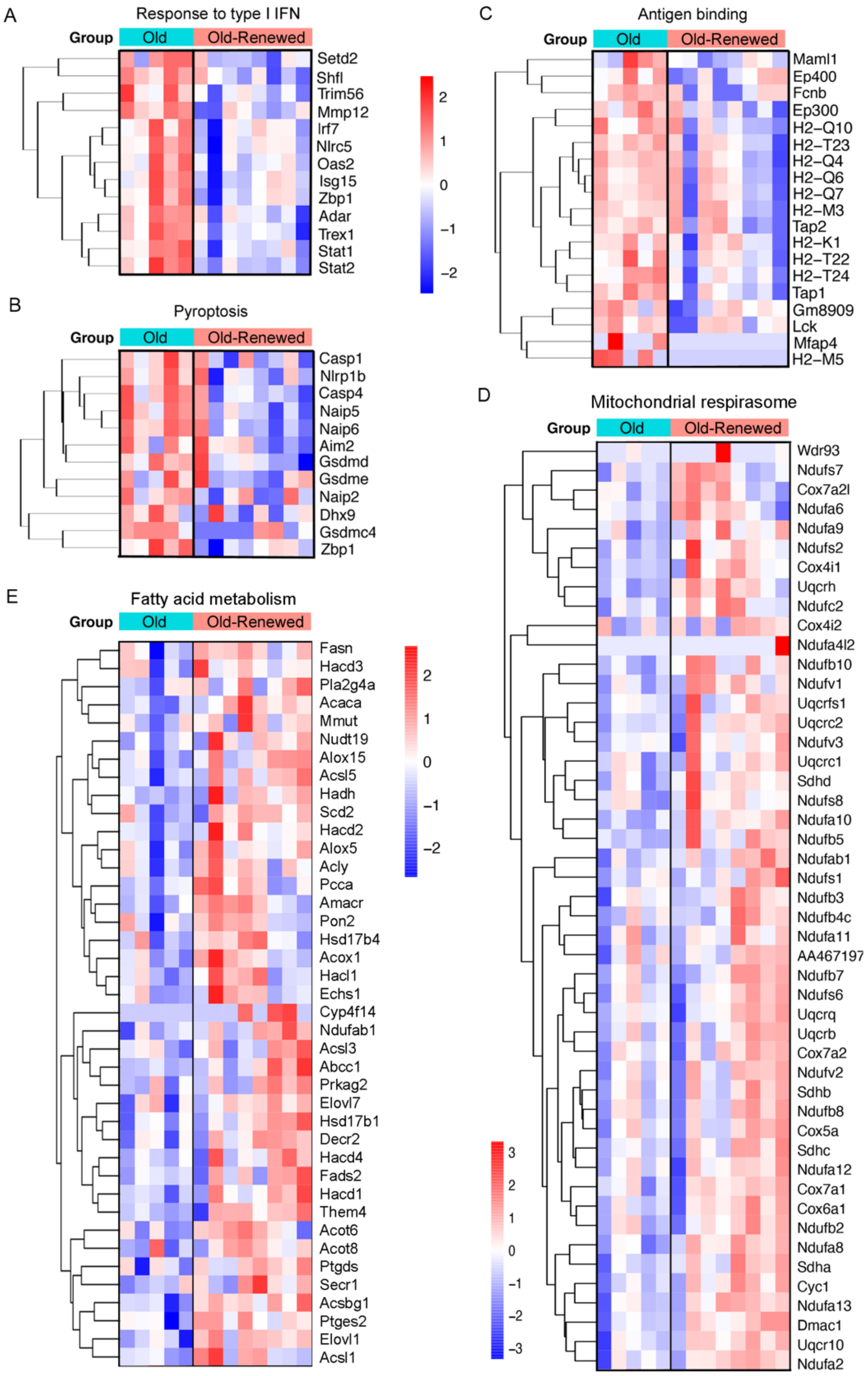


**Suppl. Fig. S4. Enrichment of genes in GO pathways in renewed microglia of old ischemic mice versus the original microglia of old ischemic mice.** *Related to Fig. 4.* RNAseq analysis of microglia obtained by FACS from the brain of old mice with repopulated microglia (Old-renewed) (n=8) versus old mice with the original microglia (n=5) four days after ischemia. Heatmaps illustrate DEGs in various GO pathways downregulated (A-C) and upregulated (D, E) in renewed microglia of old mice after ischemia in relation to the original microglia of old ischemic mice. Downregulated GO terms include: A) 'Response to type I IFN', B) 'Pyroptosis' and C) 'Antigen binding'. In contrast, enriched pathways in renewed microglia include: D) 'Mitochondrial respirasome' and E) 'Fatty acid metabolism'.

Supplementary Fig. S5

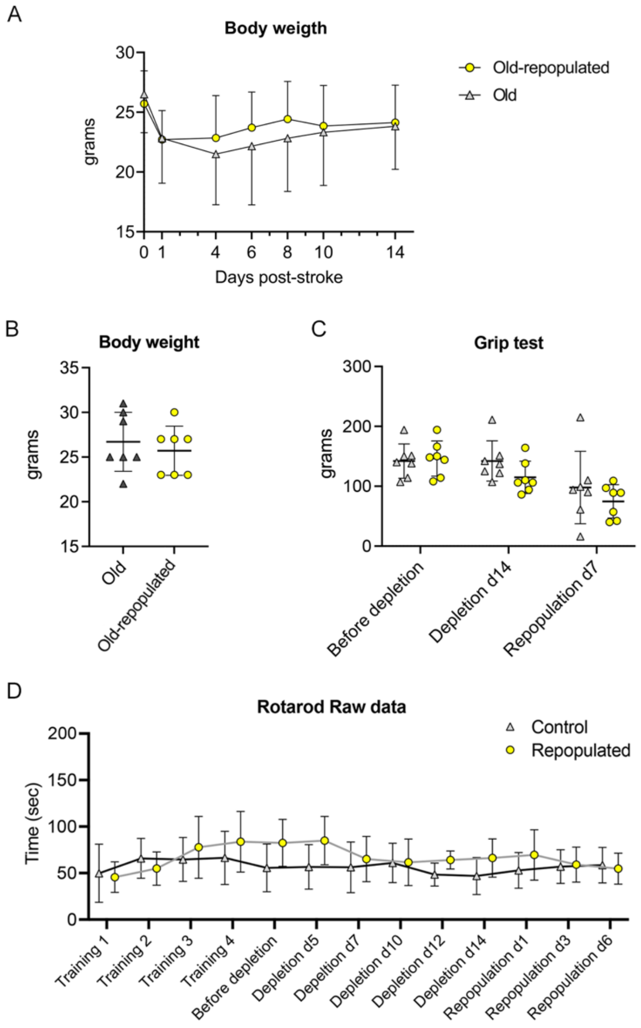


**Suppl. Fig. S5. Behavioral analysis of old mice with or without microglia depletion/repopulation**. *Related to Fig. 6.* Old female mice received PLX5622 diet for two weeks for microglia depletion followed by control diet for another week for microglia repopulation before induction of ischemia (n=7). Treatment controls received the corresponding control diet all the time (n=7). Mice were studied for 2 weeks more after ischemia. A) Mean body weight evolution after induction of ischemia in both groups shows no differences. B) Body weight after microglia repopulation (prior to ischemia) in old mice was similar in both groups regardless of the diet, as shown for each individual mouse. One mouse of the control group died after day 1 post-ischemia and was excluded from A and B graphs. C, D) Behavioral tests performed in these mice prior to ischemia showed no significant differences between groups for the grip (C) and rotarod (D) tests during treatments (two-way ANOVA, treatment effect $p=0.30$ and $p=0.34$, respectively). Values show data for individual mice and/or show the mean±SD.
